# Single nucleotide polymorphisms and structural variants reveal complex and variable ploidy in the amoebozoan *Acanthamoeba castellanii*

**DOI:** 10.1101/2025.04.25.650682

**Authors:** Morgan J. Colp, John M. Archibald

## Abstract

*Acanthamoeba castellanii* is a free-living amoeba that is emerging as a model organism for the study of eukaryotic microbiology. It is one of the most widely studied members of the Amoebozoa, and is both an important grazer in soil communities and an opportunistic human pathogen; *A. castellanii* is thus of evolutionary, ecological, and biomedical significance. Despite its potential as a lab workhorse, the genome biology of *A. castellanii* is complex and poorly understood. Polyploidy is a common feature of many amoebozoan genomes, and members of the genus *Acanthamoeba* are no exception; they appear to be not only polyploid, where genome copy number is inflated beyond the conventional haploid and diploid states, but also aneuploid, i.e., with inter-chromosomal copy number variation. To better understand aneuploidy in *A. castellanii* and how it may vary over time and between closely related strains, we analyzed nanopore and Illumina sequence datasets from several wild-type and mutant *A. castellanii* lines, with a focus on quantifying single nucleotide polymorphism (SNP) and structural variant allele frequencies across chromosome-scale scaffolds. Our findings suggest that intragenomic chromosome copy number is highly variable in *Acanthamoeba* and can change dynamically even over laboratory time scales.

**Significance Statement:** *Acanthamoeba castellanii* is becoming an important model organism for basic and applied research. However, its apparent polyploidy and aneuploidy has the potential to complicate the interpretation of results that depend on knowledge of gene copy number. In this study, we reveal the complex nature of ploidy in this organism by analyzing long- and short-read sequence data. Our results provide a reference point against which genomic and experimental data from *A. castellanii* can be interpreted, and guide future efforts aimed at more precisely characterizing how the organism regulates its genome and chromosome copy number.

## Introduction

*Acanthamoeba castellanii* is an attractive model organism for molecular and cell biological experimentation. It is easy to grow axenically to high density, it is large enough to facilitate light microscopic observation, and its cell biology is considered representative of a large fraction of eukaryotic biodiversity. At the same time, however, aspects of the nuclear biology of *A. castellanii* are complex and poorly understood, most notably its chromosome number and ploidy. In 1972, Pussard attempted to use cytological methods to count chromosomes in *A. castellanii*, but found they were too small and tightly clustered to do this accurately (Pussard 1972). By measuring nuclear DNA content, Byers (1986) inferred the now- widely-cited estimate of 25*n* for *A. castellanii*’s ploidy. Pulsed-field gel electrophoresis (PFGE) has also been used to explore *A. castellanii* karyotypes. Rimm et al. (1988) resolved 17 chromosome-sized bands for the Neff strain, recognizing that this was likely an underestimate; the intensity of some bands were suggestive of co-migrating chromosomes, and Southern blot experiments detected nuclear genes that had not migrated out of the well, suggesting that the largest chromosomes were not resolved. Matsunaga et al. (1998) used PFGE to compare the karyotypes of 28 *Acanthamoeba* strains across 12 species and showed them to be heterogeneous, even within species. Based on variable band intensities, the authors proposed that *Acanthamoeba* genomes are aneuploid.

A draft genome assembly for *A. castellanii* was published in 2013 (Clarke et al. 2013), but while useful from a gene discovery perspective, it was produced using Illumina short-read technology and thus highly fragmented. More recently, a combination of nanopore long-read and Hi-C technology allowed near-complete resolution of all *Acanthamoeba* chromosomes into scaffolds (Matthey-Doret et al. 2022), which makes it possible to revisit the question of ploidy in this organism. These assembled chromosomes serve as a predicted karyotype to a level of detail that has thus far been impossible in PFGE experiments, while the amplification-free library preparation method allows relative quantitation of the chromosomes; the absolute per-cell counts cannot be determined from these data, but ratios of sequence depth of coverage can hint at relative abundances of different chromosomes in the presence of aneuploidy. With high throughput sequencing, data on allele frequencies across chromosomes can also be collected, bringing additional, distinct lines of evidence to the question. Long read data provides two independent types of allele frequency data, the more common single nucleotide polymorphism (SNP) allele frequencies and structural variant allele frequencies. Short read data provide an additional set of SNP frequencies for cross-referencing. Here we sought to shed light on the genome organization and ploidy of *Acanthamoeba* using the rich sequence data sets underlying the long-read, chromosome-scale assemblies of the wild-type Neff and C3 strains (Matthey-Doret et al. 2022) and transformed clones sequenced to study the fate of artificial transgenes (Colp et al. 2025).

## Methods

### Inferring Single Nucleotide Polymorphism Allele Frequencies from Long- and Short-read Sequence Data

We analyzed nanopore sequence data generated by two recent genomic studies on *A. castellanii* (Matthey-Doret et al. 2022; Colp et al. 2025). This includes multiple nanopore read sets of the wild-type *A. castellanii* Neff strain and a read set from the C3 strain. Sequence data from one Neff clonal isolate transformed with circular pGAPDH-EGFP, and three clonal isolates transformed with linearized pGAPDH-EGFP were also analyzed.

Wild-type Neff nanopore reads were mapped against the entire wild-type Neff assembly with minimap2 (Li 2018) v2.24 and allele frequencies of SNPs were plotted using ploidyNGS (Augusto Corrêa dos Santos et al. 2017) v3.0.0. The assembly was then divided into its individual scaffolds, and the wild-type nanopore reads generated in the Archibald Lab were mapped against each of the 35 largest scaffolds independently. PloidyNGS plots were then generated for each scaffold. Read mapping and plotting were performed as above.

To explore the stability of the observed patterns over laboratory timescales (i.e., between wild-type Neff and transformed clonal isolates) and evolutionary timescales (between Neff and C3), nanopore reads from ‘Clone 1’, ‘Clone LT6’, ‘Clone LT8’, ‘Clone LT9’, and the Institut Pasteur wild-type Neff culture were also mapped against the separated wild-type Neff scaffolds and ploidyNGS plots were generated for each scaffold using the reads from each of these data sets. The reads from the Institut Pasteur C3 culture were mapped against the C3 wild-type genome assembly instead, but the 35 largest scaffolds were still analyzed independently.

Illumina short reads were previously generated for a subset of the isolates mentioned above for the purposes of polishing nanopore data (Matthey-Doret et al. 2022; Colp et al. 2025). This includes Illumina reads from the Archibald Lab and the Institut Pasteur for the wild-type Neff culture, and from the Institut Pasteur for the C3 culture. Among the transformed cultures,

‘Clone LT6’ and ‘Clone LT9’ were sequenced with Illumina. For these isolates, Illumina reads were mapped against the reference genome using HISAT2 (Kim et al. 2019) v2.2.1 and SNP allele frequencies were plotted with ploidyNGS v3.0.0. As before, all read sets were mapped against the wild-type Neff assembly, except for C3 reads which were mapped against the C3 assembly.

### Structural Variant Calling and Allele Frequency Plotting from Nanopore Reads

The structural variant caller Sniffles 2 (Smolka et al. 2024) was used to detect structural variants in the nanopore read mapping data described above. Sniffles was configured to detect insertions or deletions of at least 50 bp. Sniffles plot was used to visually summarize variant calling data, including plotting allele frequencies of structural variants.

## Results

Using the *Acanthamoeba castellanii* strain Neff assembly (Matthey-Doret et al. 2022) ploidy was inferred using ploidyNGS (Augusto Corrêa dos Santos et al. 2017) v3.0.0. This program plots a histogram of the absolute counts for each possible allele frequency (in steps of 0.01) detected in sequencing read sets mapped to a reference genome, and the overall shape of the histogram can be used to infer ploidy. In this case, however, a very high level of background noise was observed across the possible range of allele frequencies and there were no clear peaks (Fig. S1). Given the possibility that this complex pattern was due to aneuploidy, the analysis was repeated by dividing the assembly into its constituent scaffolds and performing the same mapping and plotting analyses for each scaffold individually. This approach produced interpretable but variable signals across the chromosome-sized scaffolds (Fig. 1).

**Figure 1.**
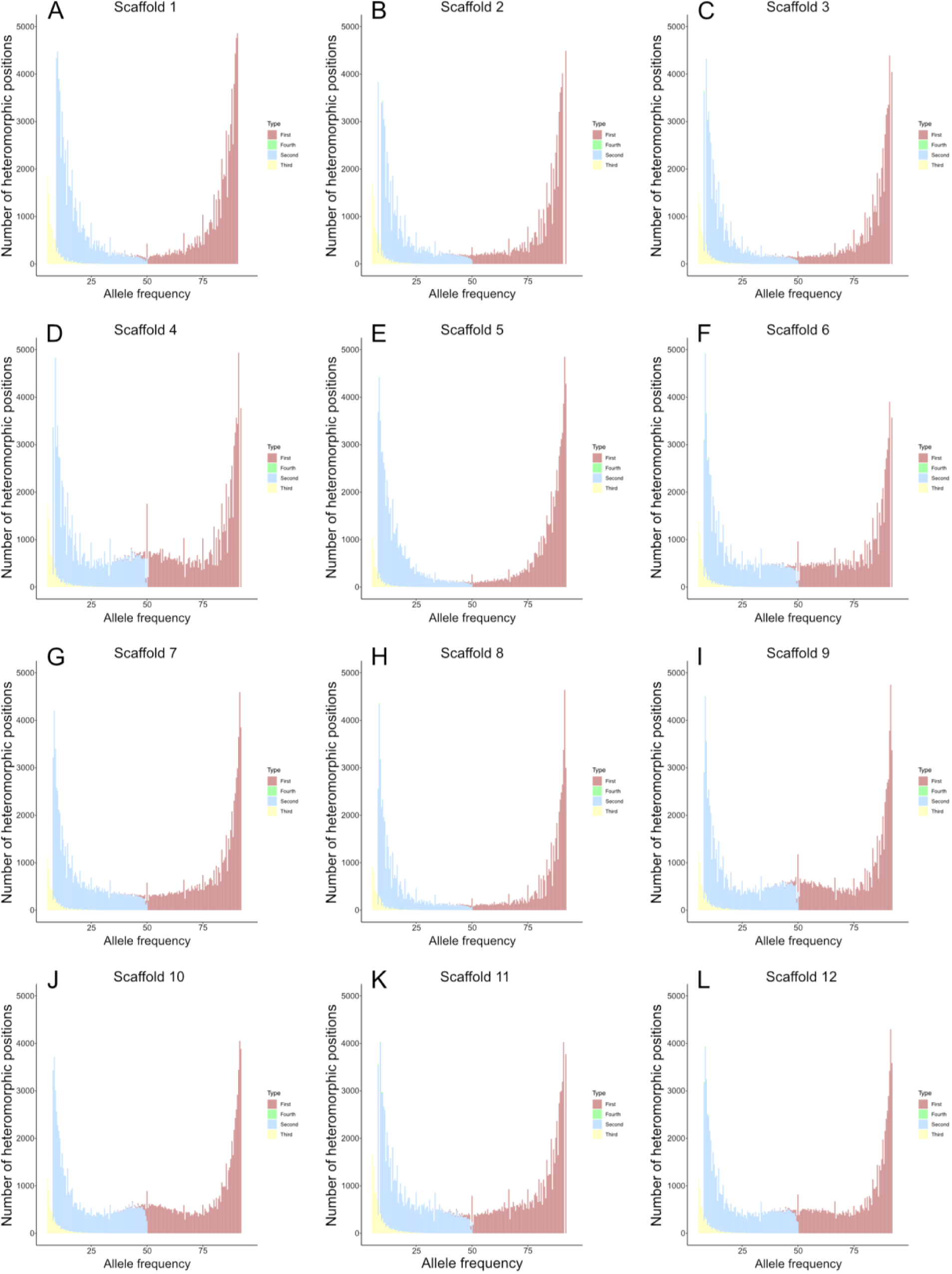
PloidyNGS SNP frequency plots from the 12 longest scaffolds of *Acanthamoeba castellanii* strain Neff. Plots were generated using nanopore reads mapped to individual reference genome scaffolds with minimap2. The ploidy signal is clearly varied across these 12 scaffolds from the same strain. A - L. Scaffolds 1 to 12, respectively.

Interpretation of these data requires a distinction between ‘ploidy signal’ and actual ploidy. Here ploidy signal refers to the lowest ploidy level that would be consistent with the observed data. For example, peaks at allele frequencies 0, 0.5, and 1 are consistent with diploidy, but it is formally possible that two identical sets of five chromosomes (10 in total) could yield this same signal. This is important because we do not have absolute quantitation of any of the *A. castellanii* chromosomes and must default to identifying the lowest ploidy level consistent with the plots as the ‘ploidy signal’, while recognizing the possibility that the true copy number is a multiple of the base number. This is also why the ploidy levels inferred here are lower than those of Byers (1986) who inferred a global ploidy of 25*n* from nuclear DNA content.

Figure 1 shows ploidy signal across the 12 largest scaffolds of the wild-type Neff assembly (ranging from 1.47 to 2.54 Mbp in size). Real data do not always cleanly conform to the above-mentioned patterns expected for different ploidy levels, but most of these plots can be interpreted. For example, the plots for scaffolds 1, 2, and 3 (Fig. 1A-C) appear to represent haploid signal, with peaks near the extremes of the plot but not between them. The peaks for Scaffold 1 (Fig 1A) do tail out further toward the middle of the allele frequency range, but no distinct peaks can be observed along those curves that would change the interpretation. For Scaffolds 5, 7, and 8 (Fig. 1E, G, H), a haploid signal is also apparent; again, the peaks at the extremes of some of the plots do taper off slowly but there are no distinct ‘shoulders’ along their curve that could be indicative of another set of allele frequencies.

The plots for Scaffolds 4, 9, 10, and 12 (Fig. 1D, I, J, L) all closely resemble what one would expect for a diploid chromosome, with peaks at the extremes and a smooth peak centering around an allele frequency of 0.5. The plots for Scaffolds 6 and 11 are more difficult to interpret (Fig. 1F and K). Here there are allele frequencies represented somewhere between the two peaks at the extremes, but these additional peaks are not well resolved. For Scaffold 11, the peaks from the extremes of the plot tail down toward the centre, and there is a small inflection in the corresponding curve around allele frequencies 0.33 and 0.67 which could represent a triploid signal. The plot for Scaffold 6 is the least clear; here there could be additional peaks near 0.33 and 0.67 once again, or potentially 0.4 and 0.6 which may correspond to pentaploidy. Regardless of the exact interpretation of these data, these examples illustrate how these plots can be used to infer ploidy signal in general and across different scaffolds within a genome, which is consistent with aneuploidy in *Acanthamoeba*.

Having found support in the long-read sequencing data for aneuploidy in the wild-type *A. castellanii* Neff genome, other independent read sets from different cultures were analyzed in the same way to ask several related evolutionary questions. These additional cultures included (i) another wild-type culture of Neff, maintained and sequenced separately by collaborators at the Institut Pasteur, (ii) a wild-type culture of *A. castellanii* C3, also maintained and sequenced at the Institut Pasteur, and (iii) four monoclonal isolates of Neff derived from wild-type cultures in the Archibald Lab at Dalhousie University, transformed by the same plasmid, and then individually bottlenecked into clonal cultures. Three of these were transformed with linearized pGAPDH-EGFP in the same experiment, while one was transformed with a circular form of the same plasmid in a separate experiment (Colp et al. 2025). Interestingly, the results showed variation in the inferred ploidy profiles across all chromosome-level scaffolds within and between the different datasets.

The transformed clonal isolates were found to vary in ploidy signal among one another, as well as compared to the wild-type culture from which they were derived. In the case of Scaffold 10, for example, while the two independently maintained Neff cultures have the same diploid signal (Fig. 2A and B), the signal in strain C3 (Fig. 2C) appears more likely to be triploid. The four clonal isolates, derived from wild-type Neff in the Archibald Lab, show additional variation for this scaffold. Tetraploid signal for Scaffold 10 is indicated by peaks at allele frequencies 0.25 and 0.75 in Clones LT8 and LT9 (Fig. 2F and G), while triploid signal appears in Clone LT6 (Fig. 2E). At first glance, the signal in ‘Clone 1’ (Fig. 2D) appears triploid, but closer inspection reveals peaks that are closer to allele frequencies of 0.4 and 0.6, corresponding to a pentaploid signal. Not all scaffolds show this degree of inter-strain signal heterogeneity, but Scaffold 10 is emblematic of the broader patterns observed, particularly in and among the clonal isolates when compared to the wild-type cultures.

**Figure 2.**
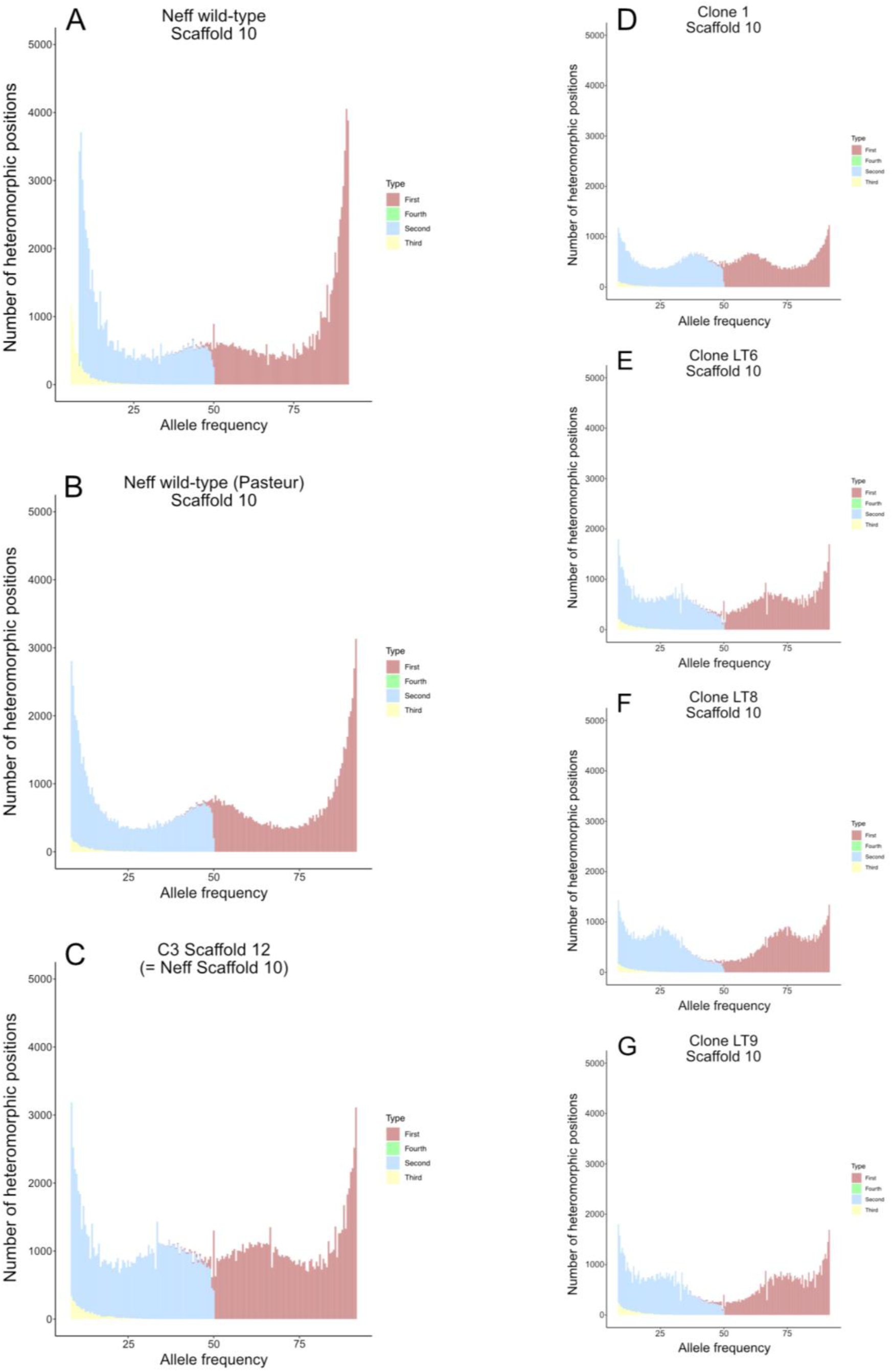
PloidyNGS SNP frequency plots for Scaffold 10 (12) across seven *Acanthamoeba castellanii* isolates. Plots were generated from long reads mapped to the Neff reference genome with minimap2. A. Scaffold 10 with reads mapped from the wild-type culture of the Neff strain in the Archibald lab. B. Scaffold 10 with reads mapped from the wild-type culture of the Neff strain sequenced at the Institut Pasteur. C. Scaffold 12 from the C3 strain, which is homologous to Scaffold 10 in Neff, with reads mapped from C3. D. Scaffold 10 with reads mapped from Clone 1. E. Scaffold 10 with reads mapped from Clone LT6. F. Scaffold 10 with reads mapped from Clone LT8. G. Scaffold 10 with reads mapped from Clone LT9.

The two wild-type Neff datasets were also found to differ in inferred ploidy signal for several chromosomes, despite both having been acquired first-hand from the American Type Culture Collection under the same strain ID by the respective laboratories at Dalhousie and the Institut Pasteur prior to sequencing. It is worth noting that neither were bottlenecked into monoclonal isolates prior to genome sequencing. Finally, variation in ploidy signal between the two wild-type Neff cultures and the C3 strain were observed, with the C3 strain being more different from either Neff culture than they were from each other. Evolutionary time and distance does appear to correlate with the degree of variation in effective ploidy signal, but this correlation is not easily quantifiable.

Two additional lines of evidence support the allele frequency-based inferences from the long read data: Illumina-based ploidyNGS allele frequency plots and structural variant allele frequency information from long read data. When visualizing long reads mapped to the *A. castellanii* reference genome assemblies, clear instances of structural variation could be observed in the mapped reads and coverage plots, where subsets of reads were missing segments of DNA ranging from a hundred to a few thousand base pairs, with an associated, cleanly delineated drop in depth of coverage. Importantly, structural variants on the same chromosome were found to share the same allele frequency (where alleles correspond to the presence or absence of the indel) when they came from the same dataset, suggesting that at least some of the signal is biological.

The long read structural variant calling program Sniffles 2 (Smolka et al. 2024) was used to make structural variant calls on all chromosomes independently, using each distinct long read dataset, such that each SNP-based ploidyNGS plot across the range of chromosomes and datasets had a corresponding structural variant-based Sniffles 2 histogram of allele frequencies (Fig. 3).

**Figure 3.**
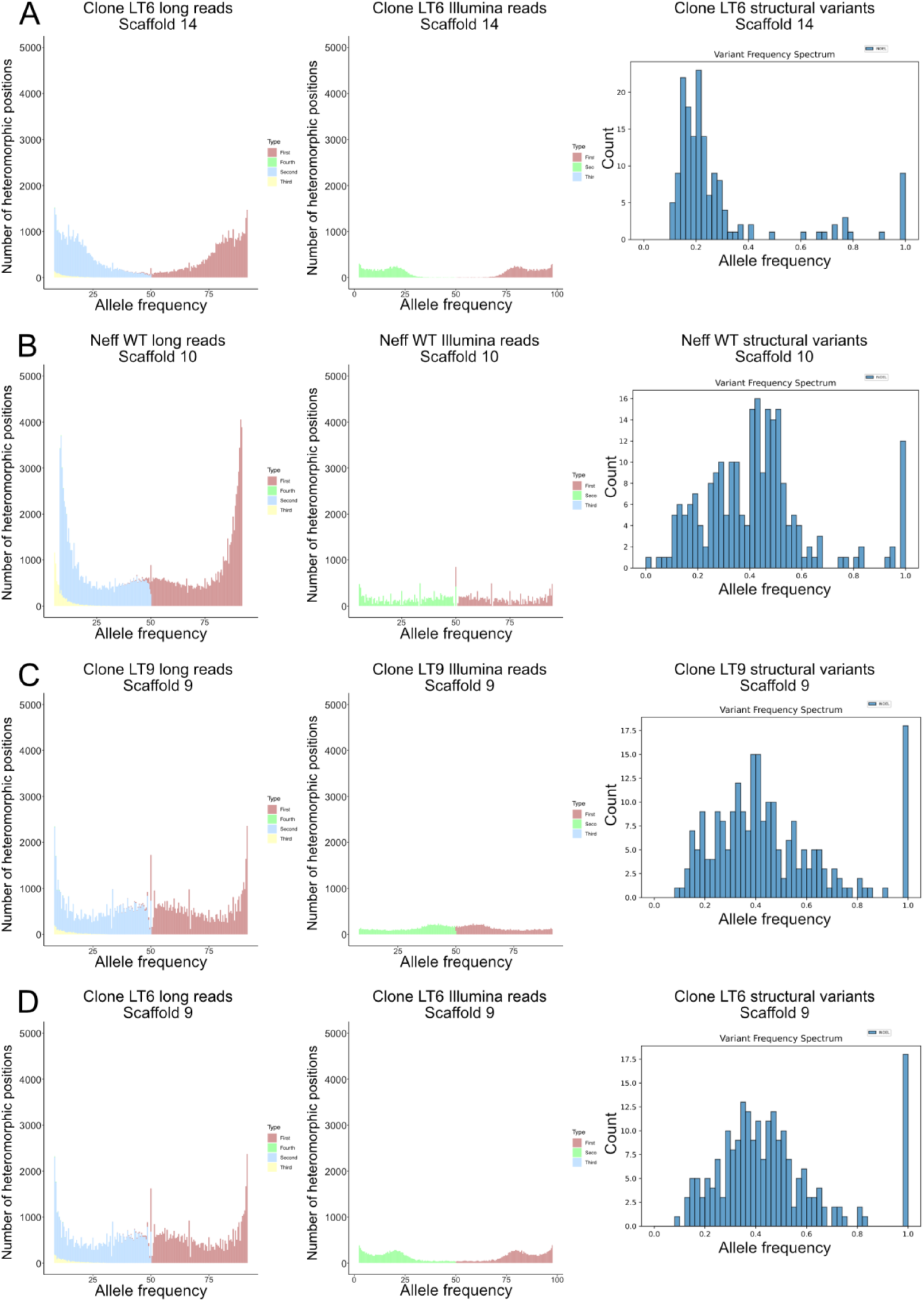
SNP and structural variant allele frequency plots for four *A. castellanii* strain Neff isolate-scaffold combinations. A. PloidyNGS plots using long and short reads from Neff Clone LT6 mapped to Scaffold 14, and a structural variant allele frequency plot generated from the same mapped long reads. B. PloidyNGS plots using long and short reads from the Archibald Lab wild-type Neff culture mapped to Scaffold 10, and a structural variant allele frequency plot generated from the same mapped long reads. C. PloidyNGS plots using long and short reads from Clone LT9 long reads mapped to Scaffold 9, and a structural variant allele frequency plot generated from the same mapped long reads. D. PloidyNGS plots using long and short reads from Clone LT6 mapped to Scaffold 9, and a structural variant allele frequency plot generated from the same mapped long reads.

To generate the ploidyNGS plots using Illumina data, the same general approach was used as for the long reads, with appropriate adjustments, such as the specific choice of read mapping program. For a subset of the isolates, this meant the possibility of comparing three different ploidy signal estimates to arrive at a consensus best inference for each chromosome in each of those isolates. Given the noise and/or uncertainty in some of these plots by themselves, having multiple independent lines of evidence proved valuable. Figure 3 illustrates four different isolate-scaffold pairs for which all three types of data are available, albeit with differing degrees of agreement between the three.

The first two examples, from Scaffold 14 in *A. castellanii* transformant Clone LT6 (Fig. 3A) and Scaffold 10 in the wild-type Neff culture from the Archibald Lab (Fig. 3B), are cases where inference was already straightforward; here the three types of data support our initial inferences. In the first case, peaks around allele frequencies of 0.2 and 0.8 are observed from the SNP frequency data (Fig. 3A), with the more accurate Illumina data providing much tidier plots. The structural variant allele frequency plot (Fig. 3A, far right) is consistent with both SNP-based datasets, with a peak around an allele frequency of 0.2 for the variant. These three plots all converge on an inferred pentaploid signal for Scaffold 14 in Clone LT6. In the next set of plots, Scaffold 10 from the Archibald Lab Neff wild-type culture (Fig. 3B), the SNP frequency plot for the long-read data has a well-defined peak at allele frequency 0.5. The plot for the Illumina data is not as clean but is consistent with this. The structural variant plot also has its peak centered around allele frequency 0.5; again, the three data sets agree on a diploid signal for Scaffold 10 in the wild-type Neff culture maintained in the Archibald Lab.

The third example, Clone LT9 Scaffold 9 (Fig. 3C), is one where one or more of the plots would have been misleading or difficult to interpret in isolation, but the combination of the three allows an inference to be made. Here the SNP frequency plot from the long-read data roughly resembles a diploid signal with a peak at an allele frequency of 0.5, but with a dip directly in the centre of the peak. The Illumina SNP frequency plot shows peaks on either side of 0.5, probably at allele frequencies of 0.4 and 0.6, and the structural variant allele frequency plot has a peak centered around 0.4, which confirms the apparent signal from the Illumina SNP frequency plot. Therefore, combining the information from all three plots reveals that the signal for Scaffold 9 from strain Neff Clone LT9 is probably pentaploid.

Finally, we identified instances in which the long-read, short-read, and indel data do not agree. With Neff Clone LT6 Scaffold 9 (Fig 3D), for example, the long-read SNP frequency plot looks like that of Clone LT9 Scaffold 9, with a peak centered around 0.5 with a small dip in the middle. The structural variant allele frequency plot, also like LT9 Scaffold 9, is more consistent with peaks at allele frequencies of 0.4 and 0.6. However, the Illumina SNP frequency plot clearly shows peaks at 0.2 and 0.8. It is worth noting that both 0.4/0.6 and 0.2/0.8 peaks are consistent with a pentaploid signal, but the ratios of one haplotype to the other are different between the two samples.

Based on these three different lines of evidence, inferences of the ploidy signal were compiled for the 30 largest chromosomes across all *A. castellanii* Neff isolates, recognizing that some had more information available than others. Only the top 30 scaffolds were considered because for the five smallest chromosome-scale scaffolds, the plots were insufficient to allow meaningful inferences. For the C3 strain, Circos plot data generated previously (Matthey-Doret et al. 2022) were used to determine which scaffolds were homologous to the top 30 Neff scaffolds. In many cases this could not be determined unambiguously, due to apparent or suspected chromosomal rearrangements disrupting chromosome-wide synteny. The data were nevertheless recorded and in cases where there were clear discrepancies between the long and short reads, ploidy levels inferred from long read sequence data were prioritized. The goal was not to precisely define the ploidy of each chromosome across all isolates, but to assess the *variability* in ploidy signal across chromosomes and isolates. The ploidy signals were inferred and recorded based on all available information but not necessarily unambiguously recorded for each plot individually; this is because cross-referencing of signals from two or three plots was sometimes required to make sense of the data. As shown in Figure 4, marked variation in ploidy signal can be seen across scaffolds within a strain and between homologous scaffolds across the studied strains.

**Figure 4.**
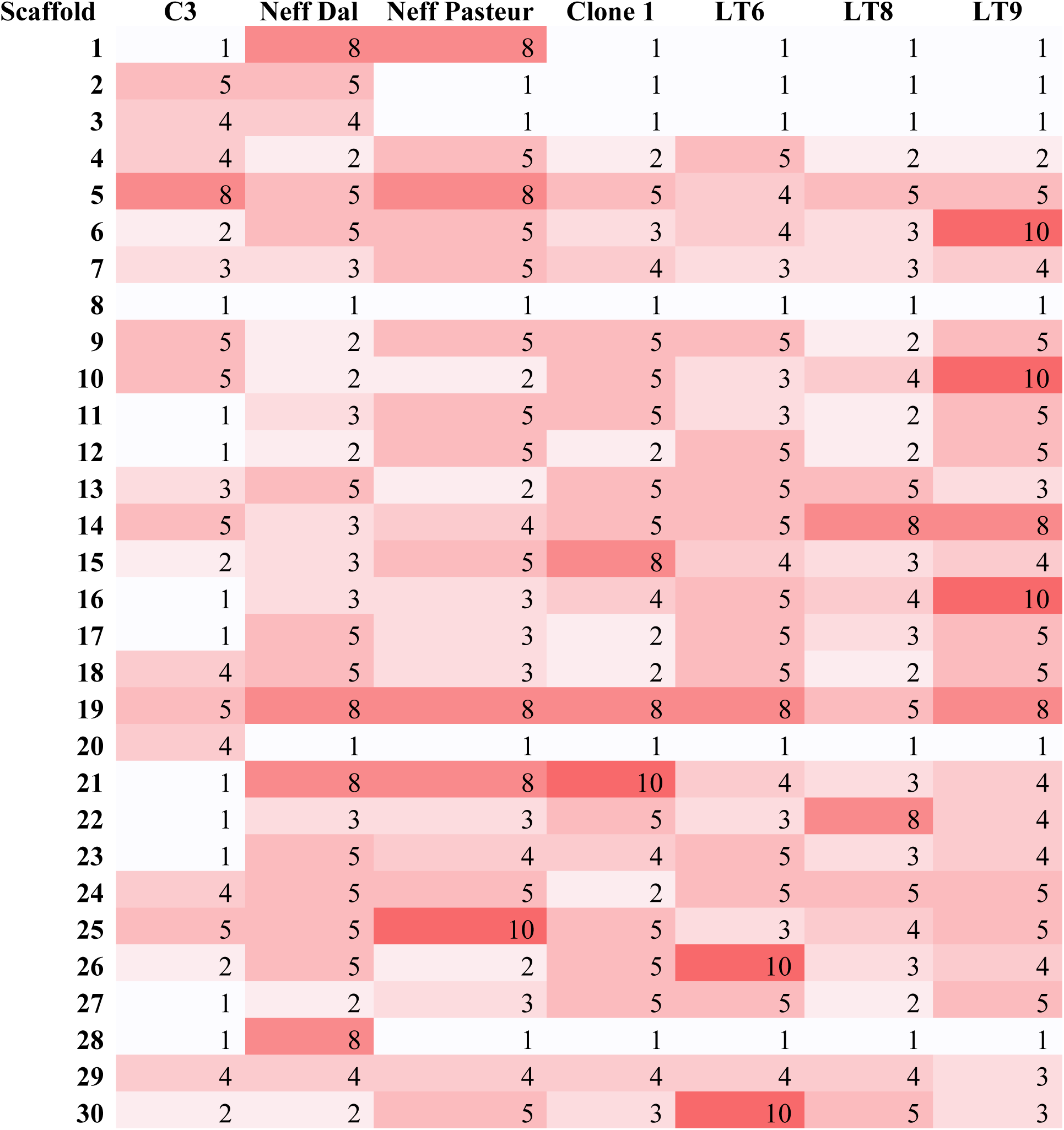
Inferred ploidy signal for the top 30 scaffolds of six *Acanthamoeba castellanii* strain Neff isolates and homologous scaffolds in *A. castellanii* strain C3. Inferences are based on the aggregate of information in ploidyNGS plots using both nanopore and Illumina sequence reads and structural variant allele frequency plots using nanopore reads. The colour scale ranges from lowest (white) to highest (red) inferred ploidy signal.

## Discussion

The goal of this study was to use a wealth of new sequence data from multiple *Acanthamoeba castellanii* isolates to improve our understanding of the organism’s ploidy. Our allele frequency-based results do not provide absolute quantification of the chromosomal copy number, but they do reveal patterns of variation in effective ploidy signal across isolates, as well as between chromosomes within a given isolate. This information helps provide bounds on the range of possible ploidy signals for individual *A. castellanii* chromosomes, as well as insight into how such levels can vary over laboratory and evolutionary timescales.

The SNP- and indel-based approach employed herein was not without limitations.

Resolving peaks on the SNP frequency plots proved challenging, and as the signal in the data moves toward a higher ploidy level, it becomes difficult to distinguish heterozygous alleles with a 1:*n*-1 ratio from homozygous alleles. As a result, it can be difficult to make precise inferences at any ploidy level. Even apparently haploid and diploid signals, which one might expect to be easiest to identify, require careful assessment. This is exemplified in Figure 3C (Scaffold 9 of *A. castellanii* strain Neff, Clone LT9) in the case of apparent diploid signal, and as described below for haploidy. Ploidy levels may also be systematically underestimated because without a high degree of structure in the allele frequency plots, estimating the signal will tend toward less granular interpretations of exactly where peaks appear. For example, without extremely narrow peaks, an actual allele frequency of 0.083, consistent with alleles in a 1:11 ratio across 12 chromosome copies, could be largely indistinguishable from an allele frequency of 0.1, consistent with alleles in a 1:9 ratio across 10 chromosome copies. In this case, one would likely tend toward the more ‘conservative’ inference of 10 chromosome copies. Another issue is that the three wild-type datasets are not from clonal isolates, so inferences may be less precise due to mixed signals in the data from diverging clones within the same culture. Importantly, each of the transformed Neff cultures analyzed herein was established from a single cell (Colp et al. 2025) and the data from these clonal isolates still suggest a complex ploidy state, supporting the idea that it is not simply an artefact of within-culture clonal heterogeneity. Even though we cannot pinpoint ploidy levels precisely, there is still value in considering overall trends in the data, and what those trends tell us about the dynamic nature of ploidy in *A. castellanii*.

The first axis to consider is variation in ploidy signal across chromosomes within the same isolate. We inferred as many as seven unique ploidy levels across the 30 largest scaffolds of individual isolates (Figure 4), and a lack of peak resolution may be masking even higher numbers. Inferences ranging from haploid to decaploid signal were made across all the isolates and chromosomes investigated. This speaks to the existence of considerable variability in chromosomal copy number. It is formally possible that in at least some strains all chromosomes are present in equal copy but that some are more heterogeneous than others. For example, if an isolate was found to have only haploid, diploid, and tetraploid signal across its chromosomes, it could be the case that all chromosomes are present in four copies. However, to explain all the different ploidy signals observed by that logic, the copy number would have to be divisible by all inferred ploidy levels, making it unreasonably high.

If we accept that there genuinely is a substantial amount of variation in copy number across the chromosomes of any given *A. castellanii* cell, then it is worth considering what the biological advantage(s) of such variation might be. It is possible that gene dosage plays a role, but the patterns are so complex that it is difficult to imagine cells varying the copy number of individual chromosomes for the purposes of regulating gene expression on a genome-wide scale. Additionally, the observed variation across isolates seems at odds with this hypothesis if we assume that the regulation of gene expression is stable over short evolutionary timescales; their ploidy signal clearly is not. One hypothesis for the benefits of polyploidy in asexual amoebae is that it is advantageous to combat Muller’s ratchet (Maciver 2016). However, intracellular variation in chromosomal copy number also seems unlikely to play a role if this hypothesis were true. The simplest explanation for the varied copy number is that the differential amplification of different chromosomes is stochastic and of no overarching importance. However, in this case, there must be some short- to medium-term stability within an isolate, because within clonal isolates the ploidy signal seems relatively stable.

The second axis of variation detected in this study is the change in effective ploidy signal of any given chromosome across isolates. Despite the short- to medium-term stability observed within a given isolate, there is clear variation in signal across isolates (Figure 4). This hints at the existence of ‘infrequent’ events that somehow alter or reset chromosome copy number, interspersed with periods of relative stability. At first glance, there also appears to be some correlation between evolutionary relatedness and ploidy signal, as careful inspection of the data in Figure 4 reveals that strain C3 is qualitatively more different from the Neff isolates than they are from one another. However, many of the differences involve either C3 or the different Neff isolates having a haploid signal while the other does not. Rather than reflecting a divergence in the actual maintenance of chromosome copy number, this could also be explained by a difference in heterogeneity of the chromosomes where C3 is an outlier. The haploid signal in these cases could simply result from having very few polymorphisms on a given chromosome to infer ploidy signal from, while it is more heterogeneous in the other species. It is important to recognize that stochastic amplification and bottlenecking of chromosome copy numbers could easily result in the loss of allelic diversity in these populations, which, in extreme cases, could result in homogeneous chromosome copies from which haploid signal is detected.

One explanation for the short- to medium-term ploidy stability observed within isolates but variation between them could be the existence of a loosely controlled endoreplication-type process that occurs somewhat infrequently, or, in terms relative to the generation time of *A. castellanii*, very infrequently. This process could amplify the different chromosomes to different levels, after which each one would be faithfully replicated and segregated during each round of mitosis. A hint from our data at this type of mechanism is the apparent existence of two underlying haplotypes for each chromosome. SNP frequency plots with an effective ploidy of tetraploid or higher, such as in Figure 2F, never appear with the full range of possible peaks. For example, if a chromosome was permanently pentaploid, it may be expected that there would be at least some alleles with each of the possible 1:4 and 2:3 ratios, but only one or the other is detected. Perhaps it is much more likely that a single point mutation occurs at a given locus than two of the same mutation in two copies, so a 1:4 ratio would not be particularly surprising, but we observed a non-trivial number of 2:3-dominant plots as well. This phenomenon could be explained by having a genome defaulted to effective diploidy, but when endoreplication occurs, each haplotype is differentially amplified.

Another piece of evidence in favour of the endoreplication hypothesis comes from PCR amplification, cloning, and Sanger sequencing of specific genomic loci from the wild-type Neff strain. From the sequencing of five individual clones per locus, only two major haplotypes are apparent, with only very minor additional heterogeneity being found in some clones (unpublished; Dudley Chung, pers. comm.). This would not be expected if more than two haplotypes were present at roughly equal frequency in the genome. If there truly are only two underlying haplotypes in *A. castellanii*, the proposed endoreplication process generating variation in chromosome copy number would also require a mechanism to return the genome back to its diploid state. Taking inspiration from microbial eukaryotes with characterized cycles of endoreplication and reduction, a few known mechanisms could conceivably achieve this.

Foraminifera employ a striking method known as genome segregation where cells with one highly polyploid nucleus undergo many consecutive rounds of mitosis to produce a great number of haploid gametes (Goetz et al. 2022; Timmons et al. 2024). The amoebozoan *Amoeba proteus* is also known to participate in cyclic polyploidy (Makhlin et al. 1979; Afon’kin 1986), but until recently, its approach to genome reduction was not well understood. It has now been shown that *Am. proteus* uses a phenomenon called ‘chromatin extrusion’ to reset its ploidy just prior to mitosis by expelling the excess DNA from the nucleus to be degraded (Goodkov et al. 2020).

Another amoebozoan, *Entamoeba*, undergoes endomitosis to vary the number of nuclei in a cell, and thus also vary the genome copy number (Lohia 2003).

Of these examples, the mechanisms described in foraminifera and *Entamoeba* do not seem to fit the case of *Acanthamoeba*. Gametogenesis is not known to occur in *Acanthamoeba*, and while we occasionally observe *Acanthamoeba* cells with two to four nuclei, a mitotic mechanism for cyclic polyploidy would not explain the aneuploidy as well. In contrast, the approach used by *Am. proteus* seems compatible with the hypothetical endoreplication and reduction cycle proposed here for *A. castellanii*, but if the genome does reduce to a diploid rather than a haploid state, there must be some mechanism to retain one copy of each haplotype. To our knowledge there is currently no direct evidence to support a chromatin extrusion-type reduction method in *A. castellanii* or related species.

The parasitic kinetoplastid genus *Leishmania* is an informative example of how we might think about *Acanthamoeba* aneuploidy moving forward. In *Leishmania* it is thought that aneuploidy is a constitutive feature, but the genome is fundamentally 2*n* (Sterkers et al. 2011), like what we propose here for *Acanthamoeba.* Aneuploidy in *Leishmania* is generally referred to as ‘mosaic aneuploidy’, referring to the fact that a given population can have a diversity of karyotypes represented at any given time. Some data suggest that upon founding of a population, there is an initial diversification of karyotypes followed by a shift toward a subset becoming more dominant (Black et al. 2023). There appears to be a correlation between gene expression and chromosome copy number in *Leishmania donovani*; under drug treatment or environmental stress, there appears to be an increase in copy number of the chromosomes bearing the genes that are upregulated in response to the change in conditions (Dumetz et al. 2017). Perhaps a similar process explains, at least in part, the observed aneuploidy in *A. castellanii*.

One study of aneuploidy in *L. major* used fluorescent in situ hybridization (FISH) to study the karyotype dynamics of individual cells directly (Sterkers et al. 2011). Aneuploidy was seen to be variable among clones and strains, and that even after isolating a clone, mosaicism reappeared within the population, although the ploidy state generally resembled the parent strain. If aneuploidy functions similarly in *Acanthamoeba*, this could suggest that even in the clonal isolates analyzed here, which showed a less convoluted signal than the wild-type strains that were not bottlenecked, there could be a small amount of mosaicism emerging that was not strong enough to be detected by the methods employed herein.

In their FISH study of *L. major,* Sterkers et al. (2011) also observed a high rate of asymmetric chromosome allocation during mitosis, where, for example, three copies of a given chromosome were allocated to one daughter cell, and two copies to another. They hypothesized that this resulted from poor control of DNA replication, where one chromosome might be replicated an extra time, or one may fail to be replicated, resulting in one daughter cell receiving an extra copy or lacking a copy, respectively. Such a process could explain the apparent underlying 2*n* nature of the *A. castellanii* genome, because there would only be two major haplotypes to be segregated asymmetrically. If the replication defects were infrequent, it would also explain the apparent short-term stability of the ploidy state. In *L. major,* it was generally observed that chromosomes only reached trisomy, that is, three copies of a given chromosome, with very rare tetrasomy, while in *Acanthamoeba*, higher copy numbers are inferred herein for many chromosomes. To achieve this result, presumably the asymmetric replication and segregation of chromosomes would have to persist over more generations and/or be biased toward replication of extra chromosome copies rather than failure to replicate existing ones.

Overall, the aneuploidy studies of *Leishmania* serve as a useful framework for future studies aimed at understanding *Acanthamoeba* aneuploidy. Establishing reliable FISH protocols in *Acanthamoeba* would provide a very powerful tool to make more sense of its ploidy state and its dynamic nature.

## Conclusions

We have capitalized on a wealth of sequence data for strains of *A. castellanii* to infer appreciable variation in ploidy across the chromosomes within a given clone, as well as across clones. Although we were unable to address the question of just how polyploid *Acanthamoeba* may be, we did discover what appears to be prominent and dynamic aneuploidy. There is still much to learn about aneuploidy and polyploidy in this organism, and how it manages the varying state of its ploidy, but we did find evidence suggesting an underlying diploid state of the genome where each haplotype may be differentially amplified. Comparing these observations in *Acanthamoeba* with those gleaned from *Leishmania* and other microbial eukaryotes will hopefully yield ideas for experiments aimed at testing specific hypotheses about the precise mechanisms that generate aneuploidy in this important yet enigmatic amoeba.

## Acknowledgements

The authors thank Dr. Joran Martijn for technical assistance, and Drs. Renato Augusto Corrêa dos Santos and Diego Mauricio Riaño-Pachón for advice on the use of ploidyNGS.

## Author Contributions

M.J.C. and J.M.A. designed the study. M.J.C. collected and analyzed the data. M.J.C. and J.M.A. interpreted the results. M.J.C. wrote the manuscript with revisions by J.M.A.

## Funding

This research was supported by a grant from the Gordon and Betty Moore Foundation (GBMF5782). M.J.C. was supported by graduate student scholarships from the Natural Sciences and Engineering Research Council of Canada and Dalhousie University.

## Conflict of Interest

The authors declare no competing interests.

## Data Availability

The sequencing datasets analyzed in this study can be found through SRA on NCBI. For wild-type Neff reads sequenced at Dalhousie University: SRX4620962, SRX4620963, SRX4620964, SRX4620965 with nanopore; SRX4625411 with Illumina. For wild-type Neff reads sequenced at the Institut Pasteur: SRX12218490 with nanopore; SRX12218479 with Illumina. For C3 reads: SRX12218489 with nanopore; SRX12218478 with Illumina. For Clone 1: SRX7813525. For Clone LT8: SRX27968414. For Clone LT6: SRX27968413 with nanopore, SRX27968419 with Illumina. For Clone LT9: SRX27968415 with nanopore, SRX27968420 with Illumina.

---

**Supplementary Figure S1.**
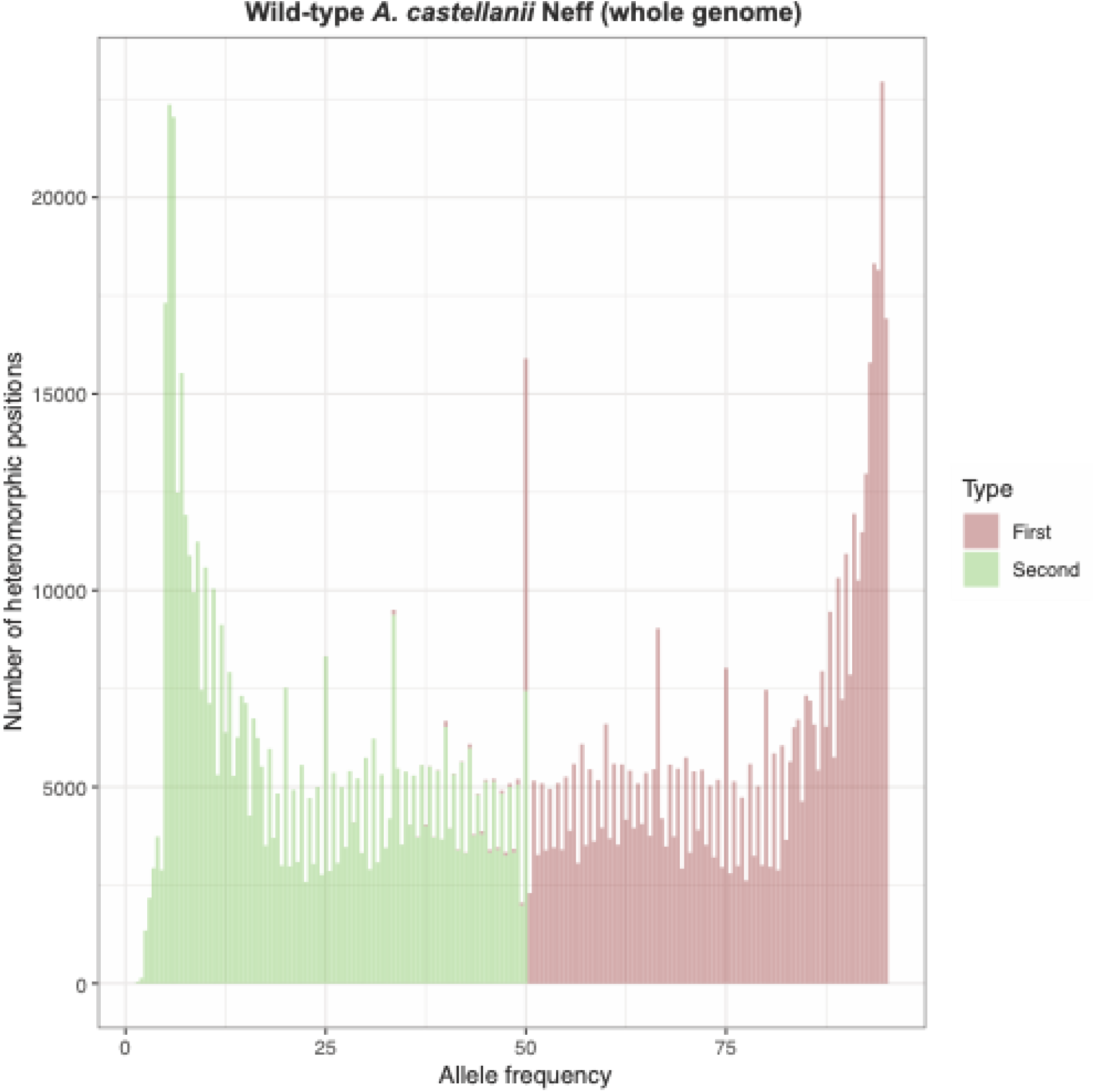
PloidyNGS SNP frequency plot of the entire *A. castellanii* Neff genome. The plot was generated using nanopore reads mapped to the wild-type Neff strain reference genome. Substantial signal can be observed across the full range of allele frequencies with no clear peaks allowing ploidy to be inferred.

## Notes

### Competing Interest Statement

The authors have declared no competing interest.

